# Meropenem vs standard of care for treatment of neonatal late onset sepsis (NeoMero1): a randomised controlled trial

**DOI:** 10.1101/456871

**Authors:** Irja Lutsar, Corine Chazallon, Ursula Trafojer, Vincent Meiffredy de Cabre, Cinzia Auriti, Chiara Bertaina, Francesca Ippolita Calo Carducci, Fuat Emre Canpolat, Susanna Esposito, Isabelle Fournier, Maarja Hallik, Paul T. Heath Frcpch, Mari-Liis Ilmoja, Elias Iosifidis, Jelena Kuznetsova, Laurence Meyer, Tuuli Metsvaht, George Mitsiakos, Zoi Dorothea Pana, Fabio Mosca, Lorenza Pugni, Emmanuel Roilides, Paolo Rossi, Kosmas Sarafidis, Laura Sanchez, Michael Sharland, Vytautas Usonis, Adilia Warris, Jean-Pierre Aboulker, Carlo Giaquinto, on behalf of NeoMero Consortium

## Abstract

**Background:** The early use of broad-spectrum antibiotics remains the cornerstone for the treatment of neonatal late onset sepsis (LOS). However, which antibiotics should be used is still debatable, as relevant studies were conducted more than 20 years ago, were single centre or country, insufficiently powered, evaluated antibiotics not in clinical use anymore and had variable inclusion/exclusion criteria and outcome measures. Moreover, antibiotic-resistant bacteria have become a major problem in many countries worldwide. We hypothesized that efficacy of meropenem as a broad spectrum antibiotic is superior to standard of care regimen (SOC) in empiric treatment of LOS and thus aimed to compare the efficacy and safety of meropenem to SOC in infants aged <90 days with LOS.

**Methods and findings:** NeoMero-1 was a randomized, open-label, phase III superiority trial conducted in 18 neonatal units in 6 countries. Infants with post-menstrual age (PMA) of ≤44 weeks with positive blood culture and one, or those with negative culture and at least with two predefined clinical and laboratory signs suggestive of LOS, or those with PMA >44 weeks meeting the Goldstein criteria of sepsis, were randomized in a 1:1 ratio to receive meropenem or SOC (ampicillin+gentamicin or cefotaxime+gentamicin) for 8-14 days. The primary outcome was treatment success (survival, no modification of allocated therapy, resolution/improvement of clinical and laboratory markers, no need of additional antibiotics and presumed/confirmed eradication of pathogens) at test-of-cure visit (TOC) in full analysis set. Stool samples were tested at baseline and day 28 for meropenem-resistant Gram-negative organisms (CRGNO).

The primary analysis was performed in all randomised patients (full analysis set) and in patients with culture confirmed LOS. Proportions of participants with successful outcome were compared by using a logistic regression model adjusted for the stratification factors.

From September 3rd 2012 to November 30th 2014, in total 136 patients in each arm were randomized; 140 (52%) were culture positive. Success at TOC was achieved in 44/136 (32%) in the meropenem arm vs. 31/135 (23%) in the SOC arm (p=0.087); 17/63 (27%) vs. 10/77 (13%) in patients with positive cultures (p=0.022). The main reason of failure was modification of allocated therapy. Adverse events occurred in 72% and serious adverse events in 17% of patients, the mortality rate was 6% with no differences between study arms. Cumulative acquisition of CRGNO by day 28 occurred in 4% in the meropenem and 12% in the SOC arm (p=0.052).

**Conclusions:** Meropenem was not superior to SOC in terms of success at TOC, short term hearing disturbances, safety or mortality and did not outselect colonization with CRGNOs. Meropenem as broad-spectrum antibiotic should be reserved for neonates who are more likely to have Gram-negative LOS, especially in NICUs where microorganisms producing ESBL and AmpC beta-lactamases are circulating.

## Intoduction

Despite significant changes in neonatal care over the last several decades, late onset bacterial sepsis (LOS) is still one of the leading causes of neonatal morbidity and mortality in developing but also in highly developed countries [1-3]. Although LOS is predominantly caused by coagulase negative staphylococci (CoNS) (36-66% of cases), Gram-negative rods are responsible for about 26-36% of cases [3, 4].

The early use of broad spectrum antibiotic regimens remains the cornerstone for the treatment of LOS. However, which antibiotic regimen should be used is still debatable, as relevant studies were conducted more than 20 years ago, were single centre or single country, insufficiently powered, evaluated antibiotics not in clinical use anymore and had variable inclusion/exclusion criteria and outcome measures [5, 6]. As a result, most antibiotics are prescribed off-label in neonates [7, 8] and treatment guidelines are based on expert opinion rather than on evidence from randomised controlled trials (RCT) [9]. As an example of this, we showed that 49 different antibiotic regimens were used for the empiric treatment of LOS in 111 patients across Europe [10]. In addition, there is significant variation in antibiotic, including meropenem, dosing in neonatal intensive care units (NICUs) [11]. The issue is now further complicated by the rise of antibiotic resistance in NICUs worldwide [12] and the paucity of new antibiotics entering the market [13-15].

Meropenem is a low protein-bound (2%), broad-spectrum carbapenem with activity against a wide variety of Gram-positive and Gram-negative bacteria including anaerobes and extended spectrum and AmpC chromosomal β-lactamase producing *Enterobacteriaceae*. Meropenem has been used off-label in NICUs for more than a decade [16] because of concerns around high rates of extended spectrum beta-lactamase producing enterobacteria and is now the second most commonly used antibiotic [11, 17]. The advantage of meropenem is its wider antibacterial coverage and thus potential of using monotherapy instead of combination therapy. However, there is serious concern around selection of carbapenem-resistant Gram-negative organisms (CRGNO)[18].

The safety and effectiveness of meropenem was recently evaluated in a single arm study including 200 infants < 91 days with suspected or confirmed intraabdominal infections. In this study, however, only 11% of patients received meropenem as monotherapy and only 15% (29/200) had positive blood cultures. The study demonstrated that meropenem was well tolerated and efficacious [19]. Meropenem was included in the European Medicines Agency priority list of off-patent drugs for which studies in neonates are requested (http://www.ema.europa.eu/docs/en_GB/document_library/Other/2013/05/WC500143379.pdf).

The general aim of the study was to suggest the appropriate use of meropenem in settings with low and medium level multi-drug resistance. Thus, the efficacy and safety of meropenem with a predefined standard of care (SOC) regimen for the treatment of LOS in patients admitted to NICU were compared. The distribution of LOS-causing microorganisms and their antibiotic susceptibility, relapse- and new infection rates, short term outcome of LOS and mucosal colonisation with CRGNO were also evaluated.

## Methods

### Study design and participants

NeoMero-1 was a randomised, open-label study conducted in 18 NICUs in Estonia, Greece, Italy, Lithuania, Spain and Turkey [20]. Patients with LOS and postnatal age (PNA) ≤ 90 days were eligible for inclusion. Culture confirmed LOS was defined as the presence of at least one positive culture from a normally sterile site together with at least one abnormal clinical or laboratory parameter within the 24 hours prior to randomisation as demonstrated in Table 1 [20]. Clinical sepsis criteria were based on postmenstrual age (PMA). If PMA was > 44 weeks the International Paediatric Sepsis Consensus Conference criteria had to be met [21]. For patients with PMA ≤ 44 weeks the criteria defined by the European Medicines Agency Expert Meeting on Neonatal and Paediatric Sepsis [5, 20] were used and the presence of at least two clinical and two laboratory parameters were required (Table 1).

**Table 1.**
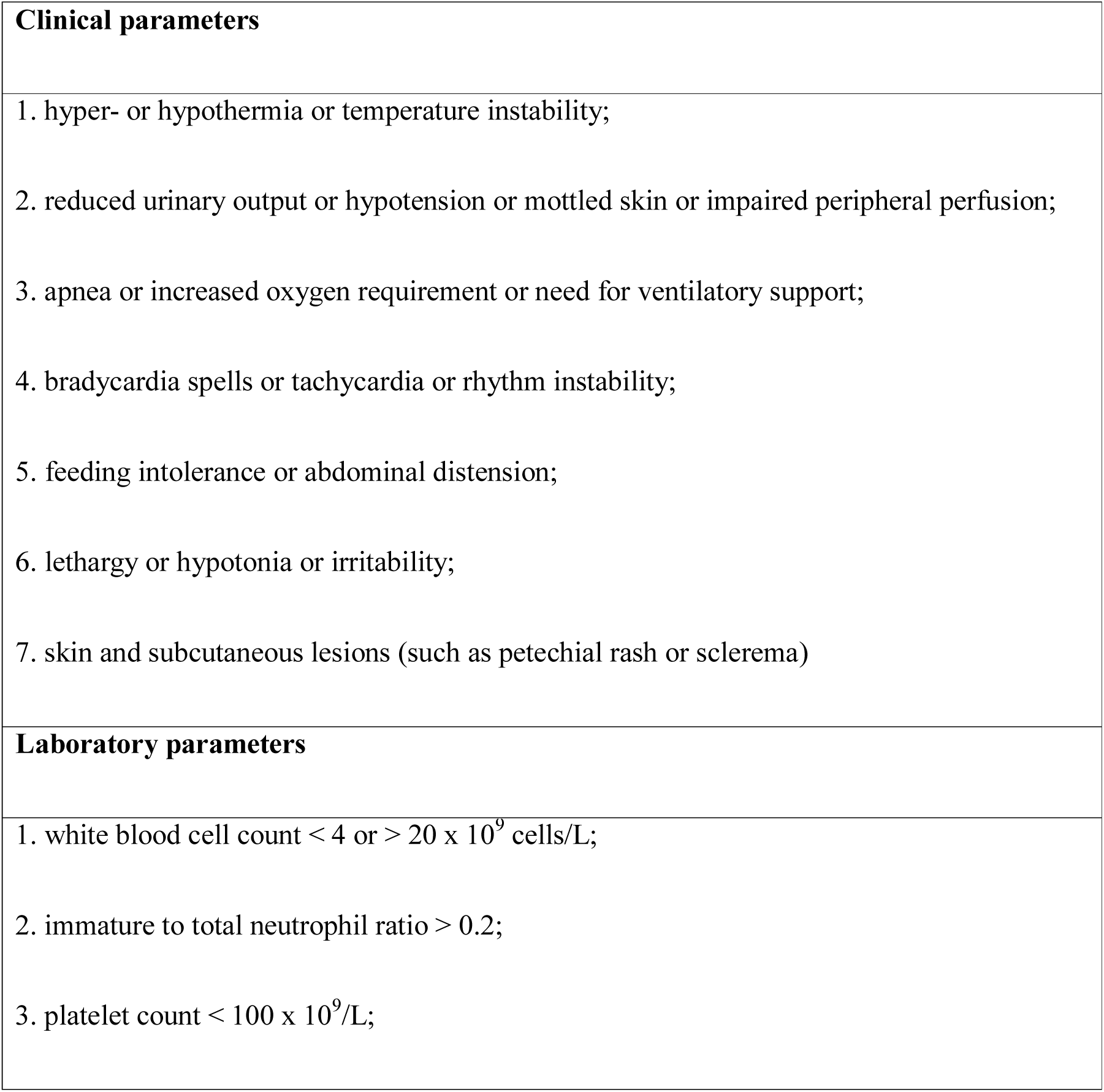
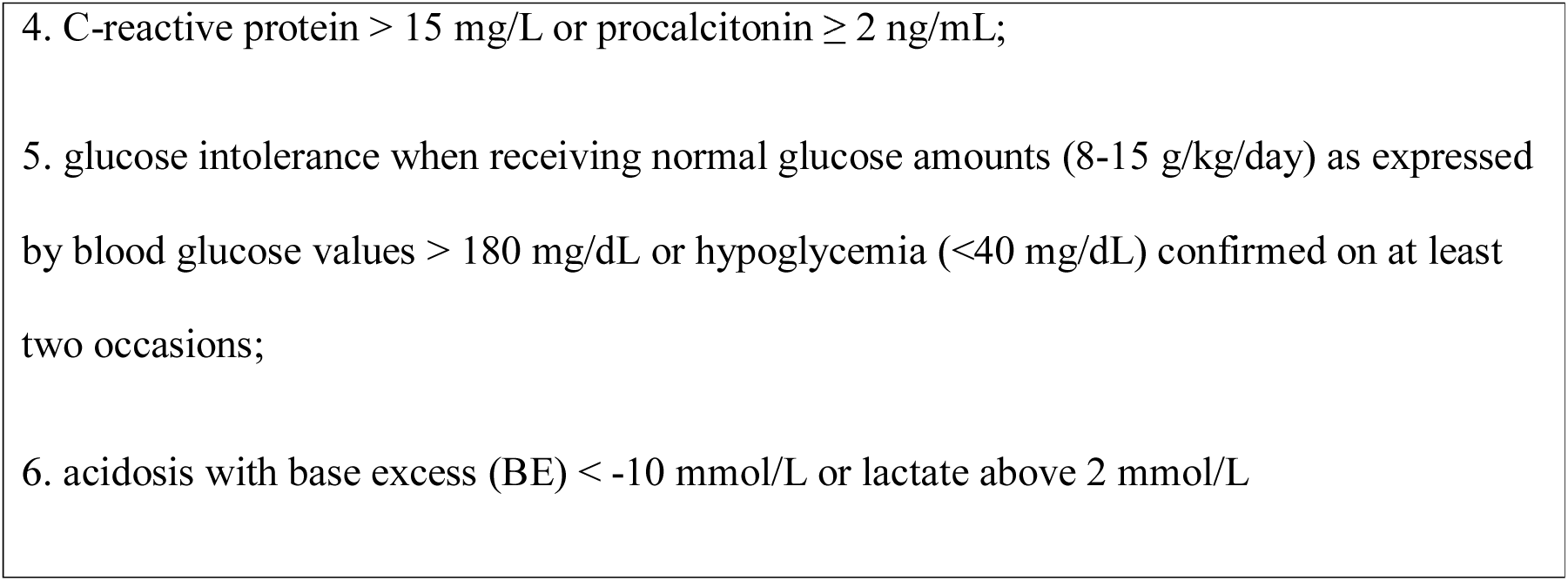
Clinical and laboratory parameters defining LOS in patients with PMA ≤ 44 weeks

Patients who had received systemic antibiotics for more than 24 hours within the 7 days prior to randomisation (except treatment failures), had meningitis and/or organisms suspected or known to be resistant to study antibiotics, were not expected to survive for more than three months, had renal failure and/or required hemofiltration or peritoneal dialysis, were excluded.

### Randomisation

Patients were centrally randomised using a computer generated randomisation list (1:1 ratio) to either meropenem or one of the two SOC regimens (ampicillin + gentamicin or cefotaxime + gentamicin) chosen by each site prior to the start of the study. Patients were stratified by SOC regimen and use of systemic antibiotics for LOS in the 24 hours prior to randomisation.

### Procedures

Meropenem was given via 30-minute intravenous infusion at a dose of 20 mg/kg q8h with the exception of those with gestational age (GA) < 32 weeks and PNA <2 weeks who received the same dose q12h with the possibility to increase dosing frequency to q8h from a PNA of two weeks. Ampicillin, cefotaxime and gentamicin were administered according to the British National Formulary for Children (BNFC, www.bnfc.org). Total duration of allocated therapy was predefined as 8 to 14 days. The concomitant use of other systemic antibiotics was not allowed with the exceptions of vancomycin, teicoplanin or linezolid, if started pre-randomisation. The use of topical anti-infectives, systemic antifungals, antivirals, immunoglobulins and probiotics was permitted.

Patients were examined at Day 0 (screening and randomisation), Day 3, end of antibacterial therapy (EOT) and test of cure (TOC) visit, which was performed 2 ± 1 days after EOT for patients treated with antibiotics for the predefined duration (11 ± 3 days). Short-term follow-up visit was performed on Day 28 by on-site visit or telephone call.

Microbiological samples were taken at baseline, Day 3, on appearance of any new signs suggestive of LOS and repeated until the relevant microorganisms were no longer detected. All samples were processed at local laboratories according to their own guidelines. In a post-hoc analysis two experts (IL and JG) reviewed susceptibility data and categorised organisms as susceptible, non-susceptible to study antibiotics, or not possible to categorise. Rectal swabs were collected within 72 hours of baseline, at EOT and at Day 28 visit or NICU discharge, and stored locally at −80°C before being periodically transferred to the central Biobank. The samples were then sent in regular batches to St George’s, University of London, Department of Medical Microbiology. The thawed faecal samples were cultured using selective media and tested for carbapenem resistance according to EUCAST guidelines (http://www.eucast.org/ast_of_bacteria/guidance_documents). The isolate was considered CRGNO if phenotypic resistance was detected to meropenem or if *Stenotrophomonas maltophilia* was isolated, and to be highly CRGNO if meropenem MIC values were ≥8 mg/L. Acquisition of CRGNO during the study was defined if these microorganisms were not detected at baseline but were found in subsequent colonisation cultures.

Hearing was assessed according to local protocol between EOT and Day 28 visit. Cerebral ultrasound (and if persistently abnormal, magnetic resonance imaging or computed tomography) was undertaken at any time between EOT and Day 28 visit.

Blood and cerebrospinal fluid samples were collected for pharmacokinetic assessment; the results of this are reported separately [22].

### Outcomes

The composite primary endpoint was assessed at the TOC visit and defined as success if (1) the patient was alive, and (2) all baseline clinical and laboratory parameters that defined LOS were resolved or improved, (3) there was no need to continue antibiotics, (4) the baseline microorganisms were eradicated or presumably eradicated with no new microorganisms identified, and (5) allocated therapy was given for 11 ± 3 days without any modification for more than 24 hours.

The secondary outcomes were safety, clinical and laboratory response on Day 3, and EOT, survival at Day 28, time to NICU discharge, presence of hearing disturbances and abnormalities in brain ultrasound, acquisition of CRGNO in rectal swabs and occurrence of relapses or new infections after successful outcome at TOC visit until Day 28. Clinical relapses were defined as recurrence of LOS together with initiation of a new course of antibiotic treatment, and microbiological relapse as an isolation of a phenotypically similar organism from a normally sterile site in a patient with signs of infection.

### Statistical analysis

On limited data available, we estimated that failure rate in the control arm would be 36% [2]. The required sample size to show a reduction of failure rate by about a third (from 36% to 23%) with 80% power in the meropenem arm using a 2-sided test at an alpha level of 0.05, was 220 patients per arm. Using a clinical definition of LOS, an ineligibility rate of 15% to 20% was anticipated. The sample size was thus conservatively increased to 275 subjects per arm to compensate for the dilution effect. Recruitment was closed on November 30th, 2014 at 272 patients randomised, due to expiration of funding by the European Commission. Considering the unexpected overall high rate of failures (70% instead of 36% due to frequent modifications of allocated therapy) and the very low percentage of subjects not having LOS, we calculated that the study had already yielded 80% power to show a 20% reduction of the failure rate, well beyond the objective of the trial.

The primary analysis included all randomised patients (full analysis set - FAS). Analysis of the primary endpoint was also performed in patients with culture confirmed LOS. Proportions of participants with successful outcome were compared by using a logistic regression model adjusted for the stratification factors. Additional efficacy analyses were performed by ignoring the changes in allocated therapy due to safety reasons or all changes of allocated therapy and by allowing duration of allocated therapy between 7 and 14 days. Other efficacy endpoints included clinical response at Day 3, end of allocated therapy and EOT, new infection and/or relapse by day 28.

Survival at day 28 was described using Kaplan-Meier method and curves were compared using a log rank test. A significance level of 5% was used and all p-values were the results of two sided tests.

All analyses were performed with the use of SAS software, version 9.3 (SAS institute).

### Ethics and registration

The local Ethics Committees approved the study protocol. The informed consent was signed by parents/guardians prior to randomisation.

The study was overseen by an independent data safety monitoring board and was registered in EudraCT database (2011-001515-31) and in clinicaltrials.gov (NCT01551394).

### Role of funding source

This study was funded by the European Commission under the FP7 program (grant number 242146) but they had no role in study design or in the analysis of data. Chiesi Farmaceutici S.P.A. provided meropenem and collaborated in the study management.

## Results

### Study population and baseline characteristics

A total of 277 infants were consented and 136 in each arm underwent randomization from September 3rd 2012 to November 30th 2014. In the SOC arm 48 (35%) patients were assigned to ampicillin + gentamicin and 88 (65%) to cefotaxime + gentamicin (Figure 1). One patient with a major informed consent violation in the SOC arm was excluded leaving 271 patients to be analysed for efficacy; 140 (52%) of them had culture proven LOS. There were 268 (99%) patients who received at least one dose of allocated therapy and were included in the safety analysis.

**Figure 1.**
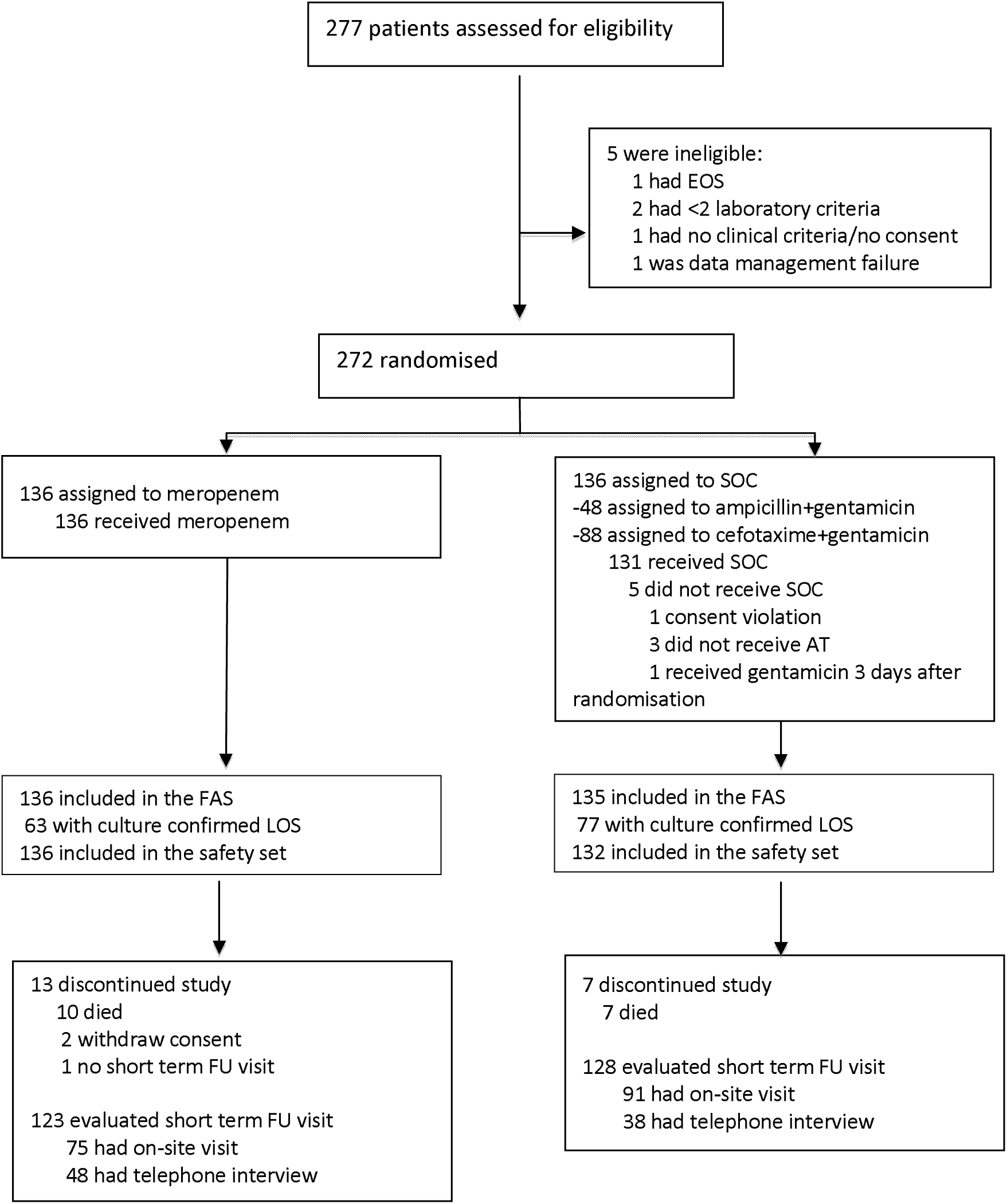
Flowchart of the study NeoMero-1. EOS – early onset sepsis; SOC – standard of care; FAS – full analysis set; AT – allocated therapy; LOS – late onset sepsis; FU – follow-up

The baseline characteristics of patients were well balanced between both arms (Table 2). They were also similar when patients were sub-grouped according to prior antibiotic treatment, culture proven LOS or presence of Gram-positive or Gram-negative LOS (data not shown). Patients in the ampicillin+gentamicin sites were more mature than those in the cefotaxime+ gentamicin sites (median PMA 39.8 vs. 32.3 weeks and median BW 2560g vs. 1105g, respectively; p < 0.0001 for both).

**Table 2.**
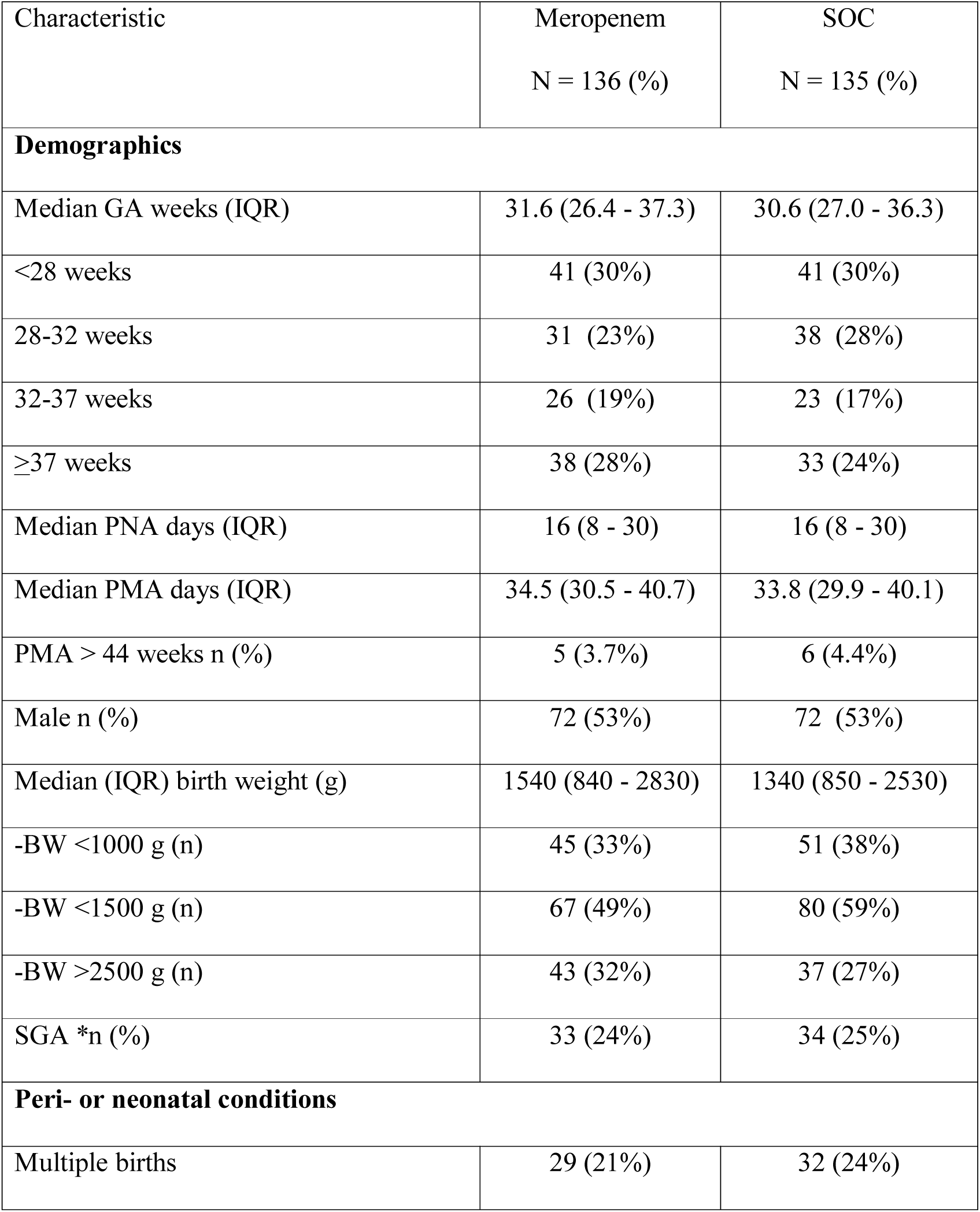
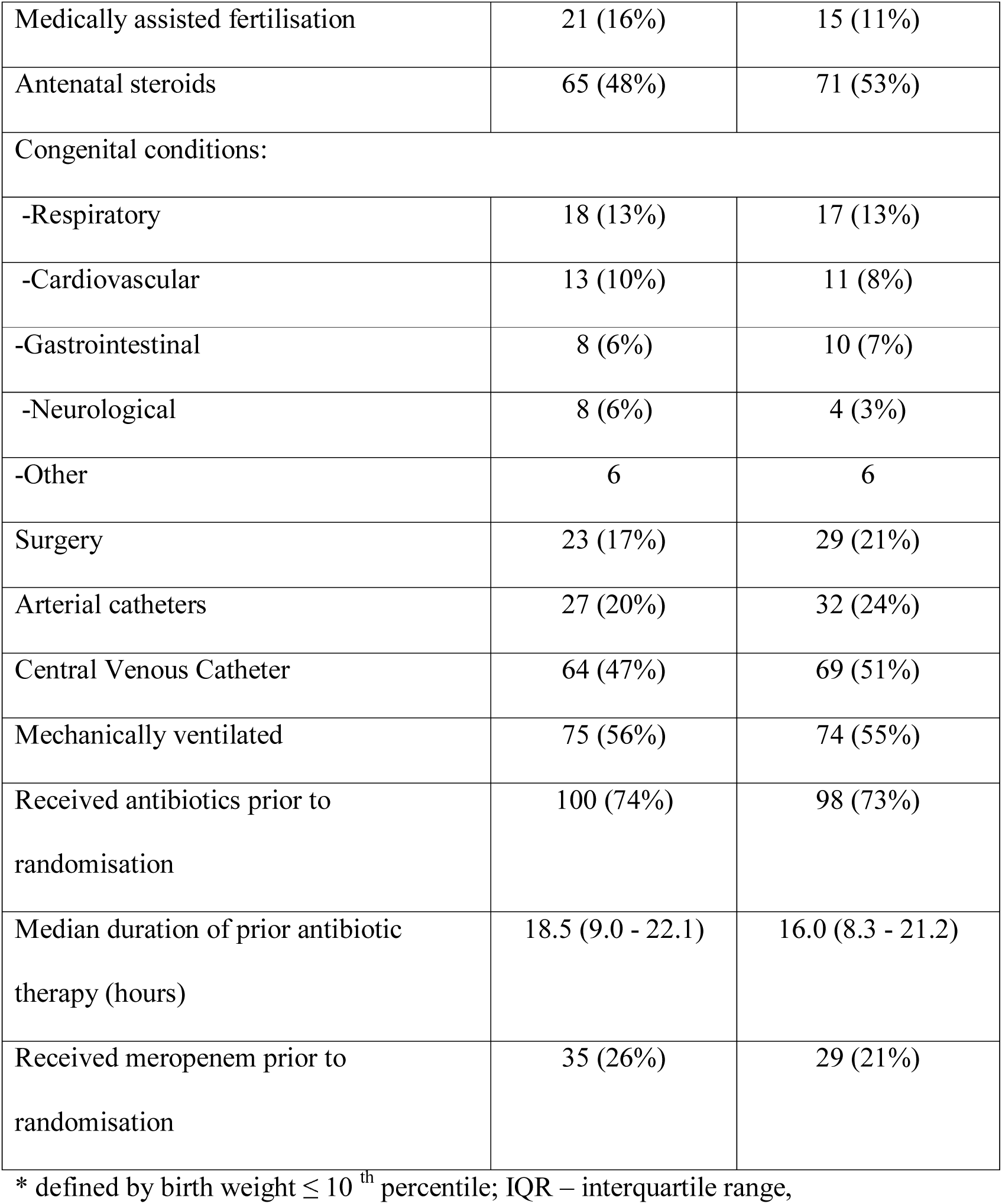
. Characteristics of study population in meropenem and SOC arm at baseline (FAS population). Data are presented as numbers (%) if not stated otherwise

In total 200 (74%) patients were premature (35% with birth weight <1000 g) and only 11 had a PMA >44 weeks. In the 24 hours prior to randomisation 73% of patients had received antibiotics; 24% had received meropenem with a similar frequency in both study arms (Table 2).

Patients of PMA ≤44 weeks had a median (IQR) of 3 (3-4) clinical and 2 (2-3) laboratory signs at baseline, in both arms. Clinical or laboratory signs seen in more than 50% of patients were impaired peripheral perfusion, mottled skin, CRP >15 mg/L and lactate >2 mmol/L (Figure 2).

**Figure 2.**
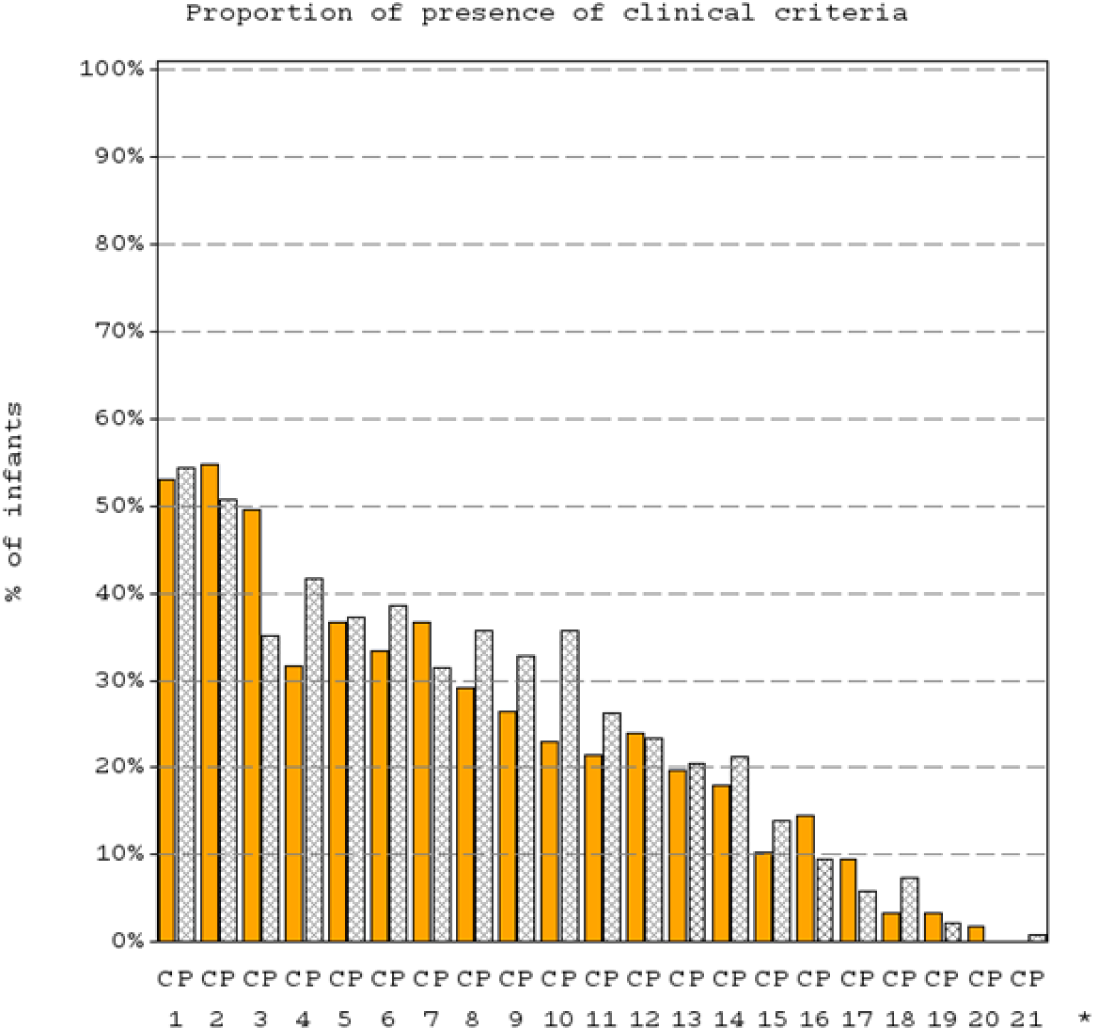
Distribution of Clinical criteria of LOS at baseline in patients of PMA < 44 weeks with clinical (C) and culture proven (P) LOS. The numbers represent the following clinical signs: **1-** Impaired peripheral perfusion, **2-** Mottled skin, **3-** Feeding intolerance, **4-**Apnoea, **5-**Increased oxygen requirement, **6-** Requirement for ventilation support, **7-** Abdominal distension, **8-** Hypotonia, **9-**Tachycardia, **10:** Lethargy, **11:** Bradycardia spells, **12:** Hyperthermia, **13:** Hypothermia, **14:** Hypotension, **15:** Other skin and subcutaneous lesions, **16:** Irritability, **17:** Rhythm instability, **18:** Reduced urinary output, **19:** T° instability, **20:** Petechial rash, **21:** Sclerema

### Aetiology of LOS

Baseline blood cultures were positive for 63/132 (46%) patients in the meropenem and 77/135 (57%) in the SOC arm with no differences in species distribution between study groups (Table 3).

**Table 3.** Causative agents of LOS and their susceptibility to study antibiotics

| Microorganism | Meropenem |  | SOC |  |
| --- | --- | --- | --- | --- |
|  | Total<br>N = 63 (%) | Susceptible to<br>meropenem<br>N (%) | Total<br>N = 77<br>(%) | Susceptible to<br>≥1 antibiotic of<br>SOC N (%) |
| **Gram-positive organisms** | **31 (49)** | **8 (26)** | **44 (57)** | **12 (27)** |
| CoNS | 22 (35) | 3 (14) | 35 (45) | 4 (11) |
| -*S. epidermidis* | 14 (22) | 2 (14) | 25 (32) | 4 (16) |
| -Other CoNS | 8 (13) | 1 (13) | 10 (13%) | 0 |
| *S. aureus* | 5 (8) | 3 (60) | 5 (6) | 5 (100) |
| -MRSA | 2 (3) | 0 | 1 (1) | 1 (100) |
| GBS | 2 (3) | 2 (100) | 3 (4) | 3 (100) |
| *Enterococcus* | 1 (2) | 0 | 1 (1) | 0 |
| Other Gram positives | 1 (2) | 0 | 0 | - |
| **Gram-negative organisms** | **24 (38)** | **22 (92)** | **25 (32)** | **18 (72)** |
| *Enterobacteriaceae* | 22 (35) | 20 (91) | 21 (27) | 16 (76) |
| - *-Enterobacter* spp. | 8 (13) | 7 (78) | 10 (13) | 6 (55) |
| - *-K. pneumoniae* | 7 (11) | 6 (86) | 4 (5) | 3 (75) |
| - *-K. oxytoca* | 4 (6) | 4 (100) | 3 (4) | 3 (100) |
| - *-Serratia spp.* | 0 | - | 1 (1) | 1 (100) |
| Non-fermentative | 2 (3) | 2 (100) | 2 (3) | 1 (50) |
| - *-Pseudomonas spp.* | 2 (3) | 2 (100) | 2 (3) | 1 (50) |
| Other Gram-negative | 0 | - | 2 (3) | 1 (50) |
| **Mixed** | **8 (13)** | **2 (25)** | **8 (10)** | **2 (25)** |

All differences non-significant between study arms; GBS – group B streptococci; MRSA – methicillin resistant *S.aureus*

All differenecs non-significant between study arms; GBS – group 318 B streptococci; MRSA – methicillin resistant *S.aureus*

Of all Gram-negative microorganisms a total of 46 (94%) were susceptible to meropenem, 17 (59%) to cefotaxime, 2 (4%) to ampicillin and 32 (65%) to gentamicin. Altogether 32/63 (51%) of all microorganisms in the meropenem and 32/77 (42%) in the SOC arms were susceptible to the allocated antibiotics.

### Antibiotic treatment

Allocated therapy was used according to the protocol in 134 (99%) of patients in the meropenem and 127 (94%) in SOC arm. In total, 65 (48%) and 67 (50%), received allocated therapy alone and 69 (51%) and 58 (43%) received concomitantly glycopeptides in the meropenem and SOC arms, respectively. The median duration of allocated therapy was comparable in both arms (7.9 [IQR 4.0-9.7] days in the meropenem vs 7.0 [IQR 2.5-9.6] days in the SOC arm; p = 0.089) but the duration of any antibiotic therapy was shorter in the meropenem than in the SOC arm (9.0 [IQR 7.8-12.0] vs 10.4 [IQR 8.5-13.3] days, respectively; p = 0.0085) (Figure 2).

### Primary efficacy analysis

In the FAS the primary outcome (i.e. the proportion of patients with a successful outcome at TOC) was comparable in both study arms - 44/136 (32%) in meropenem vs 31/135 (23%) in SOC arms (p = 0.087) (Table 4).

**Table 4.** Primary analysis: primary endpoint and culture-confirmed LOS. Data are presented as numbers (%) if not stated otherwise

|  | Primary endpoint (FAS) |  | Culture-confirmed LOS |  |
| --- | --- | --- | --- | --- |
|  | Meropenem<br>N = 136 | SOC<br>N = 135 | Meropenem<br>N = 63 | SOC<br>N = 77 |
| Treatment success at TOC | 44 (32)* | 31 (23) | 17 (27)** | 10 (13) |
| **Reasons for failure** |  |  |  |  |
| Modification of allocated therapy | 78 (57) | 85 (63) | 43 (68) | 59 (77) |
| Clinical signs not resolved or new signs | 18 (13) | 24 (18) | 8 (13) | 14 (18) |
| Microbiological failure | 3 (2) | 2 (1) | 3 (5) | 1 (1) |
| Death before TOC | 10 (7) | 6 (4) | 3 (5) | 4 (5) |
| Antibiotics not started or not-allowed antibiotics given | 2 (1) | 10 (7) | 2 (3) | 4 (5) |

\* $p=0.09$ , OR 95%CI: 1.6 (0.9 – 2.8); \*\* $p=0.02$ , OR 95% CI: 3.0 (1.2 – 7.5) (logistic model including factors of stratification)

In the culture confirmed LOS population the efficacy of meropenem was greater than that of SOC (Table 4).

The main reason for failure was modification of allocated therapy, which was more frequent in the SOC than in the meropenem arm. However, time on allocated therapy did not influence on probability of survival as shown in Figure 3.

**Figure 3.**
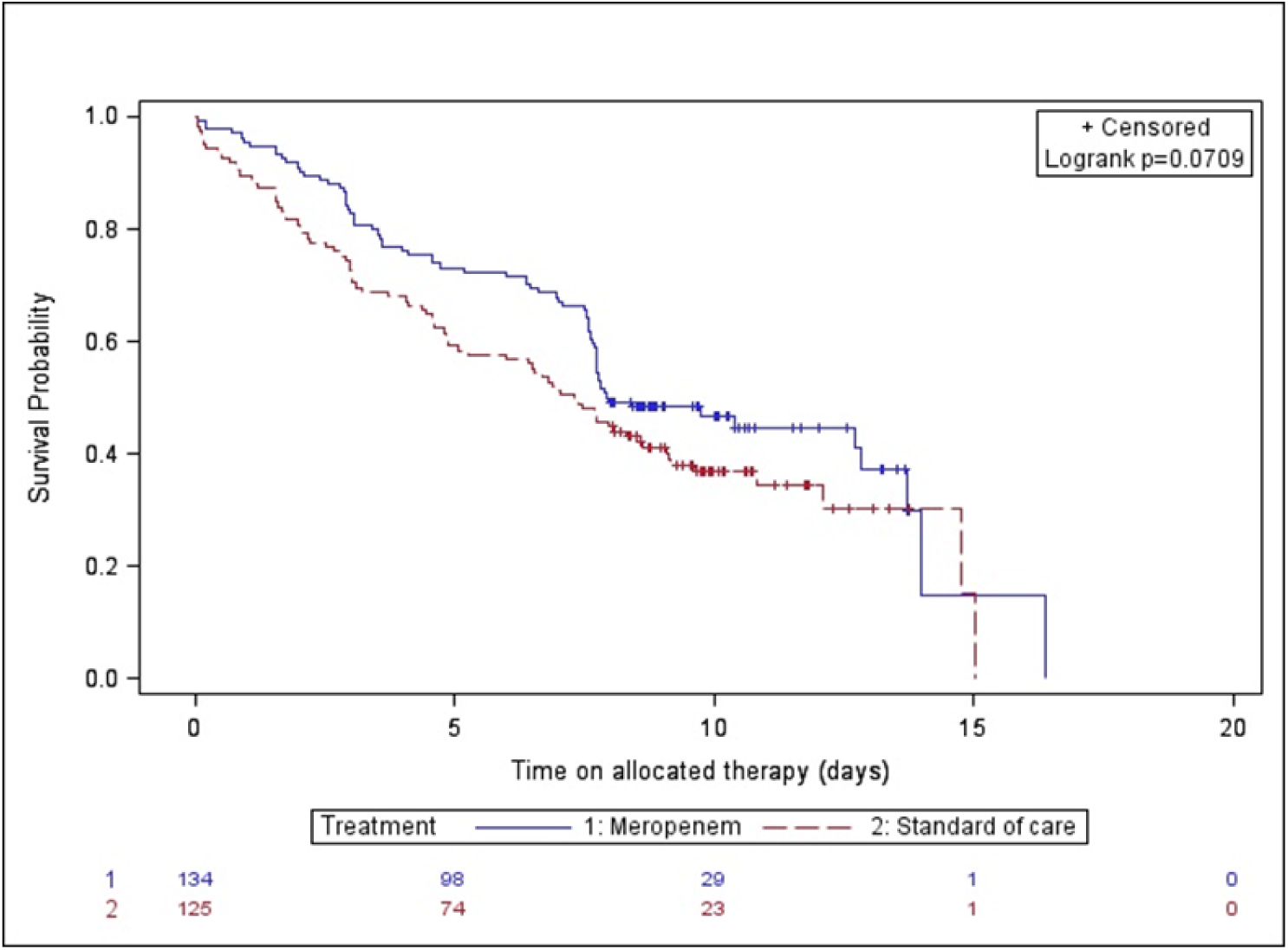
Survival probability and time to modification of allocated therapy (p = 0.0712; log-rank test). Blue indicates meropenem and red SOC

Failure was also due to completion of allocated therapy before Day 8 (38%) and diagnosis of meningitis (13%) in the meropenem arm, while isolation of resistant microorganisms (19%), lack of clinical response (18%) and inappropriate study antibiotics (18%) were the most common reasons in the SOC arm (Table 5).

**Table 5.** . Reasons for modification or discontinuation of allocated therapy

|  | Meropenem | SOC | Median duration of |
| --- | --- | --- | --- |

|  | N = 78 (%) | N = 85 (%) | allocated<br>therapy (days;<br>IQR) |
| --- | --- | --- | --- |
| Treatment completed before<br>Day 8 | 30 (38) | 10 (12) | 7.6 (7.0-7.7) |
| Meningitis diagnosed | 10 (13) | 7 (8) | 1.1 (0.2-1.7) |
| Lack of response | 8 (10) | 15 (18) | 3.1 (0.8-4.6) |
| Introduction of new and/or<br>continuation of antibiotics after<br>EOAT | 8 (10) | 5 (6) | 9.7 (8.6-12.7) |
| *Study antibiotics not needed based<br>on culture results | 5 (6) | 15 (18) | 3.0 (2.4-4.4) |
| Death | 4 (5) | 3 (4) | 1.5 (0.2-5.0) |
| Adverse event | 4 (5) | 4 (5) | 1.9 (1.3-2.7) |
| Resistant microorganism<br>isolated | 3 (4) | 16(19) | 2.9 (2.2-4.9) |
| Treatment completed after Day 14 | 1 (1) | 2 (2) | 15.0 (14.8-16.4) |
| Other | 5 (6) | 8 (9) | 4.1 (1.9-5.2) |

\*All but one patient had CoNS and 1 case had methicillin susceptible *S.aureus*

In a posthoc analysis of the FAS population, by permitting a duration of allocated therapy between 7 and 14 days (instead of 8 to 14 days), a successful outcome was more frequent in the meropenem than in the SOC arm (65/136, 48% vs 37/135, 27%; p=0.001). There were no differences in success rate between meropenem and SOC arms if changes in the allocated therapy for safety reasons were ignored (32% vs 23%) or if all changes of allocated therapy were ignored (41% vs 37%, respectively).

The success rate was greater for infants with Gram-negative than those with Gram-positive LOS (28% vs 13%; p=0.046) mainly because of the modification of allocated therapy. The success rate in Gram positive sepsis was 21% in meropenem vs 7% in SOC arm and 34% vs 23%, respectively in Gram negative sepsis; these differences were not statistically significant. The influence of vancomycin as empiric baseline therapy was tested in log-binominal model but it did not significantly influence the primary outcome.

### Secondary analysis and short term outcome

A total of 251 patients were evaluated at Day 28 either by on-site visit (66%) or by telephone interview (34%) (Figure 1). In the meropenem arm 9/61 (15%) and in the SOC arm 20/70 (29%) did not pass auditory tests (p =0.057). No differences were observed in abnormal cerebral ultrasound - 27/108 (25%) vs 30/110 (27%) in meropenem vs SOC arm, respectively. New infections or clinical relapses were seen with similar frequency in both arms (Table 6).

**Table 6.** . Secondary endpoints

|  | Meropenem<br>n/N (%) | SOC<br>n/N (%) | P value |
| --- | --- | --- | --- |
| Success at TOC based on stratification factors |  |  |  |
| No antibiotics prior to randomisation | 12/36 (33) | 7/37 (19) | 0.19 |
| At least one dose of antibiotic | 32/100 (32) | 24/98 (24) | 0.671 |
| Ampicillin+ gentamicin sites | 21/49 (43) | 18/47 (38) | 0.682 |
| Cefotaxime + gentamicin sites | 23/87 (26) | 13/88 (15) | 0.001 |
| Other factors |  |  |  |
| Patients with microorganisms susceptible to at least one component of allocated therapy | 13/32 (41) | 10/32 (31) | 0.176 |
| Alive at Day 28 | 126/136<br>(93) | 128/135 (95) | 0.462 |
| Clinical response at Day 3 | 41/125 (33) | 34/125 (27) | 0.334 |
| Clinical response at EOAT | 74/126 (59) | 60/127 (47) | 0.067 |
| Clinical response at EOT | 83/122 (68) | 76/125 (61) | 0.235 |
| New infection and/or relapse by Day 28* | 8/44 (18) | 5/31 (17) | 0.865 |

n – number of cases
N – number of patients assessed for this outcome
\*- only patients with success at TOC were evaluated for new infection/relapses

The rectal swabs were available for 130, 101 and 95 patients in the meropenem and for 127, 94, 103 patients in SOC arm at baseline, EOT and Day 28/ NICU discharge visit, respectively. Cumulative acquisition of CRGNO by Day 28 was observed in 4/94 (4%) in the meropenem and in 12/101 (12%) in the SOC arm (p = 0.052) and highly CRGNO in 3/94 (3%) and 7/100 (7%), respectively. When comparing patients who had received at least one dose of meropenem (n=170), regardless of study arm, with those not receiving meropenem, the acquisition of CRGNO in general or of highly resistant strains was similar (8/124 (6%) vs 8/71 (11%) for CRGNO and 5/124 (4%) vs 5/70 (7%) for highly CRGNO.

### Safety

A total of 193 patients (72%) had at least one adverse event (AE). All cause AEs totalled 304 and 317, with 47 and 48 serious AEs in the meropenem and SOC arms, respectively. The AEs seen in ≥3 % of patients are listed in Table 7. In the meropenem arm the most common AEs were anaemia, thrombocytopenia and meningitis and in the SOC arm anaemia, abdominal distension and apnoea. Seizures, a recognised side effect of carbapenems, were seen in four (3%) patients in the meropenem arm and one (<1%) in the SOC arm. Renal failure occurred in three (2%) patients in the meropenem arm and in four (3%) patients in the SOC arm.

**Table 7.** Comparative safety and presence of most common major clinical diagnoses in meropenem and SOC arm

|  | Meropenem<br>N = 136 (%) | SOC<br>N = 132 (%) | P |
| --- | --- | --- | --- |
| Total number of patients with AE | 91 (67) | 102 (77) | 0.059 |
| Total number of patients with grade 3/4 AEs | 51 (38) | 61 (46) | 0.148 |
| Total number of patients with SAEs | 28 (21) | 18 (14) | 0.131 |
| Discontinued treatment due to death or AEs | 8 (6) | 7 (5) | 0.796 |
| AE observed in more than 3% patients |  |  |  |
| Anaemia | 15 (11) | 24 (18) | 0.097 |
| Thrombocytopenia | 12 (9) | 5 (4) | 0.091 |
| Meningitis | 11 (8) | 5 (4) | 0.137 |
| Abdominal distension | 5 (4) | 10 (8) | 0.165 |
| Oliguria | 5 (4) | 4 (3) | 1.000 |
| Apnoea | 6 (4) | 11 (8) | 0.188 |
| Respiratory distress | 5 (4) | 3 (2) | 0.723 |
| Sepsis | 4 (3) | 7 (5) | 0.330 |
| Oxygen saturation decreased | 4 (3) | 7 (5) | 0.330 |
| Seizures | 4 (3) | 1 (1) | 0.622 |
| Hyperglycaemia | 3 (2) | 7 (5) | 0.212 |
| Major clinical diagnoses in premature neonates |  |  |  |
| RDS or HMD | 53 (39) | 62 (47) | 0.186 |
| PDA requiring surgery | 37 (27) | 37 (28) | 0.880 |
| Anaemia prematurity | 33 (24) | 37 (28) | 0.483 |
| Bronchopulmonary Dysplasia | 27 (20) | 31 (23) | 0.470 |
| Apnoea of prematurity | 24 (18) | 35 (27) | 0.080 |
| Intracranial bleeding | 21 (15) | 24 (18) | 0.548 |
| NEC stage II or worse | 11 (8) | 16 (12) | 0.273 |

Ten patients in the meropenem and seven in the SOC arm died with an overall mortality rate of 6%. While numerical differences in mortality were seen between meropenem and SOC arms in the FAS population, there were no differences in mortality in culture confirmed LOS (Table 4). The mortality rate was 1% (1/80) in Gram-positive and 10% (6/60) in Gram-negative infections. All but three patients who died had a BW <1200g.

### Discussion

We have performed the largest RCT on the efficacy of antibiotics in LOS, undertaken in a population of predominantly premature, critically ill hospitalized neonates in Europe. We have shown that the mortality was low with both antibiotic regimens and the efficacy of meropenem was similar to commonly used SOC combinations based on a complex composite primary endpoint in the FAS population. If only patients with culture proven LOS were analysed the efficacy of meropenem was significantly greater than that of SOC in general but there were no differences between study arms if Gram-positive and Gram-negative sepsis were evaluated separately. Furthermore, patients randomised to meropenem had a shorter duration of antibacterial therapy than those randomised to SOC. The two study arms were similar in terms of adverse events and acquired perirectal colonisation by CRGNO.

The NeoMero1 study differed from previous studies in LOS in many ways. First, it was a multicentre study including countries with low to moderate antibiotic resistance rates (http://ecdc.europa.eu/en/healthtopics/antimicrobial_resistance/database/) in contrast to previous single center and/or national studies [5, 19]. Second, the demanding inclusion criteria resulted in recruitment of a very sick patient population (e.g. 55% mechanically ventilated, 35% with BW of <1000g) compared to previous studies [5]. Third, only 2% of patients were ineligible (did not have LOS) and altogether 52% had culture proven LOS as opposed to 15% in a recent study of complicated intraabdominal infections [19]. Fourth, NeoMero1 had an ambitious primary endpoint that in addition to resolution or significant improvement of clinical and laboratory criteria, did not allow any changes of allocated therapy such as deviations from fixed treatment duration, dosing and/or addition of another antibiotic, in contrast to more liberal or less specific endpoints in previous studies [5, 19].

The most intriguing finding of this study, in comparison to others, was a relatively low success rate in terms of the composite primary endpoint in both study arms (23% in SOC vs. 32% in meropenem), while mortality rates were much lower than in previous studies of LOS and in a recent Egyptian study comparing conventional and prolonged infusion of meropenem [23]. The low efficacy rate was mainly driven by the modification of allocated therapy and most of all by its fixed duration of 8 to 14 days. The effect of the latter was clearly demonstrated in the post-hoc analysis in which reducing the allowed treatment duration by just one day (from 8 to 7 days) improved the success rate from 32% to 48% in the meropenem and from 23% to 27% in the SOC arms. We believe that this was due to the clinicians’ decision to stop antibiotics earlier than the pre-defined duration, presumably because they felt that clinically the sepsis episode had resolved and the infant had recovered. The optimal duration of antibiotic therapy in LOS is not known [24].

In contrast to previous studies, we did not find an association between carbapenem use and CRGNO colonization [25-27]. Of note, our study was an RCT with strict inclusion criteria, in contrast to previous retrospective and/or observational studies which included all patients without restriction [25, 27, 28]. We should emphasize that the relatively short duration (median of 9 days) of meropenem treatment in the NeoMero1 study may be relevant. For example, Clock *et al*. (2016) showed in an observational study that perirectal colonisation with Gram-negative multi-drug resistant bacteria was associated with >10 days of meropenem treatment [18].

In line with previous studies, meropenem was well tolerated and all AEs in this very sick patient population were well balanced between study arms [19]. Seizures, previously reported to be related to meropenem treatment [29], were seen in higher numbers in the meropenem arm but due to very low numbers no meaningful conclusions can be drawn.

The study had a few limitations. First, it was an open label study with the risk of investigator - induced bias when evaluating the primary endpoint or changing allocated therapy. An open label design was selected because meropenem monotherapy was to be compared with a combination of comparator agents. Using a dummy infusion in critically ill, premature babies adds significantly to the complexity and cost of a multicenter trial and is questionable from an ethical perspective. We also note that the most appropriate targets for meropenem are Gram-negative microorganisms, especially those resistant to other antibiotics like ESBL or AmpC producing organisms. Despite the demanding inclusion criteria, that well discriminated between patients with and without LOS, these criteria performed poorly in distinguishing between cases caused by Gram-positive and Gram-negative microorganisms; about half of the recruited patients still had Gram-positive infections. As long as rapid and reliable methods or biomarkers, which allow differentiation between different species, are not available, recruitment of mixed population into similar studies is unavoidable. To target antibiotic therapy more precisely, rapid and reliable tests that enable identification of microorganisms and/or their antibiotic resistance, and biomarkers that differentiate between infections and other illnesses, are urgently needed.

NeoMero1 is the first adequately powered RCT for LOS since the 1970s [5, 6] but several outstanding issues require further studies to be done. For example, the question of best treatment options for LOS in developing countries and/or in areas with high antibiotic resistance rates was not addressed as 92% of microorganisms were susceptible to meropenem and 72% at least to one component of SOC. As shown by us, RCTs in LOS treatment are challenging due to a vulnerable population and lack of validated disease criteria and endpoints [5, 6, 30]. There is an urgent need for cooperation between academia, pharmaceutical industry and regulators in innovating clinical research in neonatology, including defining alternative and more feasible study designs (e.g. pharmacokinetics/pharmacodynamics, rather than solely clinical endpoint based designs, enabling modelling/simulation and extrapolation from studies in adults) [6, 30]. It is critical to provide efficacy data for those infected with organisms covered specifically or exclusively by study antibiotics (e.g. ESBL or AmpC producing organisms).

We have also shown that the LOS criteria developed by an European Medicines Agency expert group [5] were able to discriminate well between patients with and without LOS, but further improvement and validation of these criteria is needed before adopting and implementing them into clinical trials. Indeed, other definitions have been published, which use fewer clinical and laboratory parameters, but to the best of our knowledge, these have not been tested or used in large RCTs [30]. The recent STROBE-NI consensus for reporting neonatal sepsis trials should help with this in the future [31].

## Conclusion

In predominantly premature critically ill infants with LOS in Europe, meropenem treatment was not superior to SOC in terms of success at TOC, short-term hearing disturbances, safety or mortality. However, meropenem monotherapy resulted in slightly shorter treatment duration. Meropenem did not lead to enhanced colonization with CRGNOs. We recommend that meropenem should be reserved for seriously ill premature neonates with suspected or proven Gram-negative LOS, especially in NICUs in which microorganisms producing ESBL and AmpC beta-lactamases are circulating.

## Acknowledgements

We would like to thank all patients and their parents participating in this study.

## Data safety monitoring board

Hugo Devlieger (chair), Jim Gray, John Van den Anker and Pollyanna Hardy

## NeoMero Consortium

Oguz Akbas, Antonella Allegro, Davide Bilardi, Giulia Bonatti, Nijole Drazdienė, Silvia Faggion, Eva Germovsek, Genny Gottardi, Tiziana Grossele, Cristina Haass, Tatiana Munera Huertas, Valentina Ierardi, Sandrine Kahi, Paraskevi Karagianni, Aspasia Katragkou, Eve Kaur, Birgit Kiilaspää, Karin Kipper, Aggeliki Kontou, Victoria Kougia, Hayriye Gözde, Kanmaz Kutman, Elisabetta Lolli, Valentina Montinaro, Makis Mylonas, Kader Ben Abdelkader Emmanuelle Netzer, Clarissa Oeser, Felix Omenaca, Maria Luisa Paoloni, Simona Perniciaro, Laura Picault, Carlo Pietrasanta, Andrea Ronchi, Suzan Şahin, Yacine Saidi, Marina Spinelli, Joseph Standing, Claudia Tagliabue, Tuuli Tammekunn, Nina Tiburzi

